# Comparing the Rat Grimace Scale and a composite behaviour score in rats

**DOI:** 10.1101/490482

**Authors:** Cassandra Klune, Amy Larkin, Vivian Leung, Daniel Pang

## Abstract

There is a growing interest in the use of voluntarily displayed ongoing behaviours in laboratory animals to assess the pain experience. In rats, two behavioural pain scales, the Rat Grimace Scale (RGS, a facial expression scale) and a composite behaviour score (CBS, a behavioural ethogram reliant on postural changes), are both promising pain assessment methods. Both scales have been used to assess pain in a laparotomy model, however, they have never been compared directly and the knowledge of how different analgesics may affect these two scales is limited. This study aimed to provide a comparison to discriminate the temporal and analgesic response in a laparotomy model. Female Wistar (n = 26) and Sprague Dawley rats (n = 26) were block randomized to receive saline, meloxicam (2 mg/kg) or buprenorphine (0.05 mg/kg) 30 minutes before a laparotomy model. Rats were video-recorded before surgery (BL) and at 30, 150, 270, and 390 minutes post-operatively. Videos were assessed according to both scales by a trained, blinded observer. Both CBS and RGS scores increased significantly at all post surgical timepoints in the saline group. Post-surgical CBS scores did not increase significantly above baseline levels in the groups given meloxicam or buprenorphine. However, the RGS scores only remained low in the buprenorphine group while scores increased significantly in the meloxicam group, to a similar degree as in the saline group. These findings suggest that the CBS is more sensitive to the analgesic effects of NSAIDs than the RGS.

## Introduction

Accurate and reliable pain assessment in laboratory rodents is essential to produce high quality pain research and safeguard animal welfare. The continued dependence on evoked hypersensitivity to assess pain in animals has been proposed as a contributor towards failure of translational research. While measures of evoked hypersensitivity assess hyperalgesia and allodynia, they do not capture ongoing pain which has been suggested to be most relevant in many human pain conditions [1-5]. Ongoing pain is perpetuated by an ongoing inflammatory process rather than an external stimulus. Multiple methods have been proposed to evaluate ongoing pain in animals, however, a lack of evidence describing the strengths and weaknesses of such methods in comparison to one another discourages their use [3].

From a welfare perspective, signs associated with pain are common humane endpoints in rodent research, but typical signs such as weight loss may not be specific to pain or sufficiently sensitive to be useful [6]. Traditional nociceptive tests, such as mechanical withdrawal testing, are not used as welfare assessment tools as they are time consuming and labour intensive.

Two promising behavioural assessment methods have been developed to better capture the ongoing pain experience of laboratory rodents. These are the Rat Grimace Scale (RGS) [7] and the short form Composite Behaviour Score (CBS) [8-10]. The RGS, a facial expression scale, measures the degree of change in four ‘action units’: orbital tightening, nose/cheek flattening, ear, and whisker changes. The CBS consists of counting the frequency of select full body behaviours that have been associated with pain (i.e., writhing, back arching and staggering).

Both scales are responsive to predicted changes in pain levels (increasing after a painful intervention and decreasing following analgesic administration) and have been shown to have good construct validity and inter-rater reliability [7-12]. Both scales are also relatively simple to employ. However, the independent use of these scales has provided conflicting reports on the efficacy of commonly used analgesics in reducing pain scores. CBS scores were repeatedly shown to decrease with administration of non-steroidal anti-inflammatory drugs (NSAIDs) [8-10]. However, RGS scores only decreased with particular NSAIDs or if used in very high doses [13]. This suggests that these scales may not be equally sensitive in discriminating analgesic response.

The aim of this study was to directly compare the performance of the CBS and RGS in a laparotomy model of acute post-operative pain, using meloxicam and buprenorphine, two commonly used analgesics of different drug classes. A secondary aim was to evaluate any effects of rat strain on scale by using both Wistar and Sprague-Dawley rats. It was hypothesized that scores from both scales would decrease with analgesia and that animals treated with buprenorphine would display lower pain scores compared to those treated with meloxicam. Rat strain was not hypothesized to have an effect on pain scores or analgesic response.

## Materials and methods

All experiments were approved by the University of Calgary Health Sciences Animal Care Committee, Calgary, Canada (protocol ID: AC15-0062), in accordance with Canadian Council on Animal Care guidelines.

### Animals

Adult female Wistar (n = 26) and Sprague Dawley rats (n = 27) that were at least 6 weeks of age were obtained from Charles River, Canada (250 ± 100g) or surplus stock at the University of Calgary Animal Resource Centre. Plastic cages (47 × 25 × 21cm, RC88D-UD, Alternate Design Mfg and Supply, Siloam Springs, Arizona, USA) with wood chip bedding, shredded paper, and a plastic enrichment tube were used for housing. Rats were pair housed and kept on a 12hr light dark cycle (lights on at 7:00 hours) in a temperature (23°C) and humidity (22%) controlled environment free of pathogens and negative serologically for all antigens tested on the Charles River Assessment plus. Rats were provided food (Prolab 2500 Rodent 5P14, LabDiet, PMI Nutrition International, St. Louis, MO, USA Prolab 2500 Rodent 5P14, LabDiet, PMI Nutrition International, St. Louis, MO, USA) and water *ad libitium*. Before experiments began, rats were habituated for a minimum of three days. Habituation consisted of 10 minutes of handling by each experimenter (CK and AL) and 10 minutes in a plexi-glass observation box (W 14 cm × L 26.5 cm × H 20.5 cm) in which rats would be video recorded. During handling and exposure to the observation box, rats were offered a food reward (Honey Nut Cheerios™, General Mills, Inc., Golden Valley, MN, USA). Rats were block randomized (list randomizer, random.org) to receive either meloxicam (n = 16, 2 mg/kg; Metacam Solution for Injection, Boehringer Ingelheim Vetmedica, Inc., St. Joseph, MO, USA), saline (n = 16, volume matched to meloxicam).

Following preliminary analysis, a buprenorphine treatment group was added to act as a positive drug control. A second cohort of rats were randomised to receive saline (n = 6, volume matched to meloxicam) or buprenorphine (n = 15, 0.05 mg/kg; Vetergesic, 0.3 mg/mL; Champion Alstoe, Whitby, ON, Canada). All procedures were performed during the light period, between 0730 and 1800.

### Surgery

All surgeries were performed by a single surgeon (CK). Thirty minutes before induction of anesthesia, rats received a subcutaneous injection of either saline, meloxicam, or buprenorphine. Rats were anesthetized in a plexi-glass induction chamber with 2% isoflurane carried in oxygen at 1L/min. Following loss of the righting reflex, rats were removed from the chamber and anaesthesia was maintained using a face mask. Rats were placed in dorsal recumbency on a heating pad (Sunbeam, 50watts, 120VAC, UL, USA) and ocular lubricant was placed in both eyes. The abdomen of the rat was shaved and aseptically prepared using chlorohexidine and 70% alcohol. Rats were then covered with a sterile drape so only the surgical area was exposed. A three-centimeter incision in the skin was made beginning 1 cm caudal to the xyphoid cartilage using a scalpel blade (size 15). A three-centimeter incision was made through the muscle layer by a stab incision lengthened with Metzenbaum scissors. The muscle was then closed using a simple continuous suture pattern and the skin using a subcuticular suture pattern (4-0 Monocryl (Poliglecaprone) Suture, RB-1 17mm Taper, Ethicon). Tissue glue (3M Vetbond, 3 mls, 3M Animal Care Products, St Paul, MN, USA) was used to cover knots that could not be buried in the skin. Following the skin closure, isoflurane was terminated and rats were left to recover with oxygen only. Rectal temperature was taken at this time, before the rats regained sternal recumbency. Once sternal, rats were returned to their home cages, which were warmed using a heating lamp for 150 minutes following surgery. Rectal temperatures were taken at the at 30 minute and 150 minute timepoints after video recording was completed to ensure recovery to normothermia.

### Video Recording

Rats were placed individually in the observation box in view of two cameras (Panasonic HC-V720P/PC, Panasonic Canada Inc., Mississauga, ON, Canada) positioned perpendicular to one another (one on the short side of the box and the other on the long side of the box). Rats were recorded for 15 minutes at 45 minutes before surgery (baseline; BL) and 30, 150, 270 and 390 minutes after surgery. All recordings took place in a dedicated behavioural assessment room. After the last recording period, rats were euthanized by first being anesthetised with isoflurane and, following loss of their righting reflex, overdose with carbon dioxide. Death was confirmed with cessation of heart beat.

### Video Analysis

#### The Rat Grimace Scale

The same observers took screenshots of each rats’ face throughout the 15 minute video. Four images were taken per video approximately 3.5 minutes apart but no less than one minute apart. A clear frontal view of the face was captured whenever possible. If this was not possible, a side profile was captured instead. Images were placed into presentation software (Microsoft PowerPoint, version 15.0, Microsoft Corporation, Redmond, WA, USA) and slide order was randomized (http://www.tushar-mehta.com/powerpoint/randomslideshow/index.htm). A trained observer (CK) (Zhang et al., in press) blinded to treatment and timepoint scored all images using the RGS. Action units (orbital tightening, ear changes, nose/cheek flattening and whiskers changes) were scored as either “0” if not present, ‘1’ if moderately present or “2” if obviously present, according to the original method described by Sotocinal et al. [7]. These action unit scores were averaged to produce a score between 0-2 for each image. Following unblinding of images, scores collected at each time point were averaged for each animal.

#### Composite behaviour score

Trained observers (CK and AL) blinded to treatment and time point watched the first 10 minutes of the 15 minute long video and counted the frequency of each discrete occurrence of three pain behaviours: 1) Back-arch: upward arching of the back in a cat-like behavior 2) Writhe: contraction of flank abdominal muscles producing concavity of the side of the rat caudal to the rib cage 3) Stagger: an occurrence of falling or a quicker than normal movement of the feet characterized by the loss of balance [10]. These behaviours were identified to be the major signs of pain following laparotomy by Roughan and Flecknell [10] and each occurrence of each behavior counted as one point towards the overall pain score of the rat.

### Statistical Analysis

A sample size of 12 animals per treatment group was estimated using an alpha value of 0.05 and a power of 0.8 to detect a mean difference of 0.3 with the RGS [14]. The D’Agostino and Pearson omnibus normality test was used to assess the normality of each data set. Strain differences within each treatment group were tested using two-way ANOVA with a Sidak correction for multiple comparisons. No strain differences were identified so data were pooled for further analysis. A two-way ANOVA with a Greenhouse-Geisser correction and a Tukey correction for multiple comparisons was applied to the RGS and CBS data sets to assess main effects of drug treatment and time. To assess for increases from baseline a two-way ANOVA with Dunnett post-hoc test was applied to analyse each data set. A p-value less than 0.05 was considered significant. Data are presented as mean ± SEM with the 95% confidence interval (95%CI) for the mean difference. Data were analyzed with commercial software (Prism 6.07, GraphPad Software, La Jolla, CA, USA). Data supporting the results are available in an electronic repository: https://doi.org/10.7910/DVN/CTOVDW.

## Results

Data from one rat were excluded from the buprenorphine (Sprague-Dawley) group due to a misinjection. Rats were normothermic at the time of sternal recumbency. One rat (Wistar treated with saline) dropped below the normothermic range (35.5-37.7°C, corrected rectal temperature [15]) at 30 minutes after sternal recumbency but was normothermic by the 150 minute timepoint. No other complications following surgery were encountered.

### Comparisons Between Strain

There were no differences in RGS or CBS scores between the two strains: Wistar and Sprague Dawley rats (p > 0.05; S1 table).

## Within Group Comparisons

Pain scores within the saline group increased from baseline at all post-surgical timepoints with both the RGS and the CBS (p > 0.05; Table 1, Figs 1 and 2). The RGS scores of rats treated with buprenorphine remained similar to baseline levels as did the scores from the CBS (p > 0.05; Tables 1, Figs 1 and 2). The CBS scores of rats treated with meloxicam remained similar to baseline levels at all post-surgical timepoints (p < 0.05, Fig 2). In contrast, the RGS scores for meloxicam treated rats increased significantly from baseline at all post-surgical timepoints (p < 0.05; Table 1, Fig 1).

**Table 1.** Within group comparisons (to baseline) of RGS and CBS scores at all post-surgical timespoints.

| Comparison | RGS<br>p-value [95% CI] | CBS<br>p-value [95% CI] |
| --- | --- | --- |
| **Saline** |  |  |
| BL vs. 30 | < **0.0001** [0.43 to 0.82] | **0.014** [0.30 to 3.4] |
| BL vs. 150 | < **0.0001** [0.20 to 0.59] | < **0.0001** [2.3 to 5.4] |
| BL vs. 270 | < **0.0001** [0.30 to 0.69] | < **0.0001** [2.6 to 5.7] |
| BL vs. 390 | < **0.0001** [0.24 to 0.64] | < **0.0001** [2.3 to 5.5] |
| **Meloxicam** |  |  |
| BL vs. 30 | < **0.0001** [0.55 to 1.0] | 0.90 [-1.3 to 2.3] |
| BL vs. 150 | < **0.0001** [0.18 to 0.64] | 0.39 [-0.72 to 2.8] |
| BL vs. 270 | < **0.0001** [0.31 to 0.77] | 0.29 [-0.60 to 3.0] |
| BL vs. 390 | < **0.0001** [0.25 to 0.71] | 0.69 [-1.0 to 2.5] |
| **Buprenorphine** |  |  |
| BL vs. 30 | 0.51 [-0.11 to 0.36] | 0.93 [-1.4 to 2.3] |
| BL vs. 150 | > 0.99 [-0.25 to 0.22] | > 0.99 [-1.6 to 2.0] |
| BL vs. 270 | 0.94 [-0.29 to 0.18] | > 0.99 [-1.7 to 2.0] |
| BL vs. 390 | 0.46 [-0.11 to 0.36] | > 0.99 [-1.8 to 1.8] |

p-values and 95% confidence intervals of the differences are reported for each comparison.

**Fig 1.**
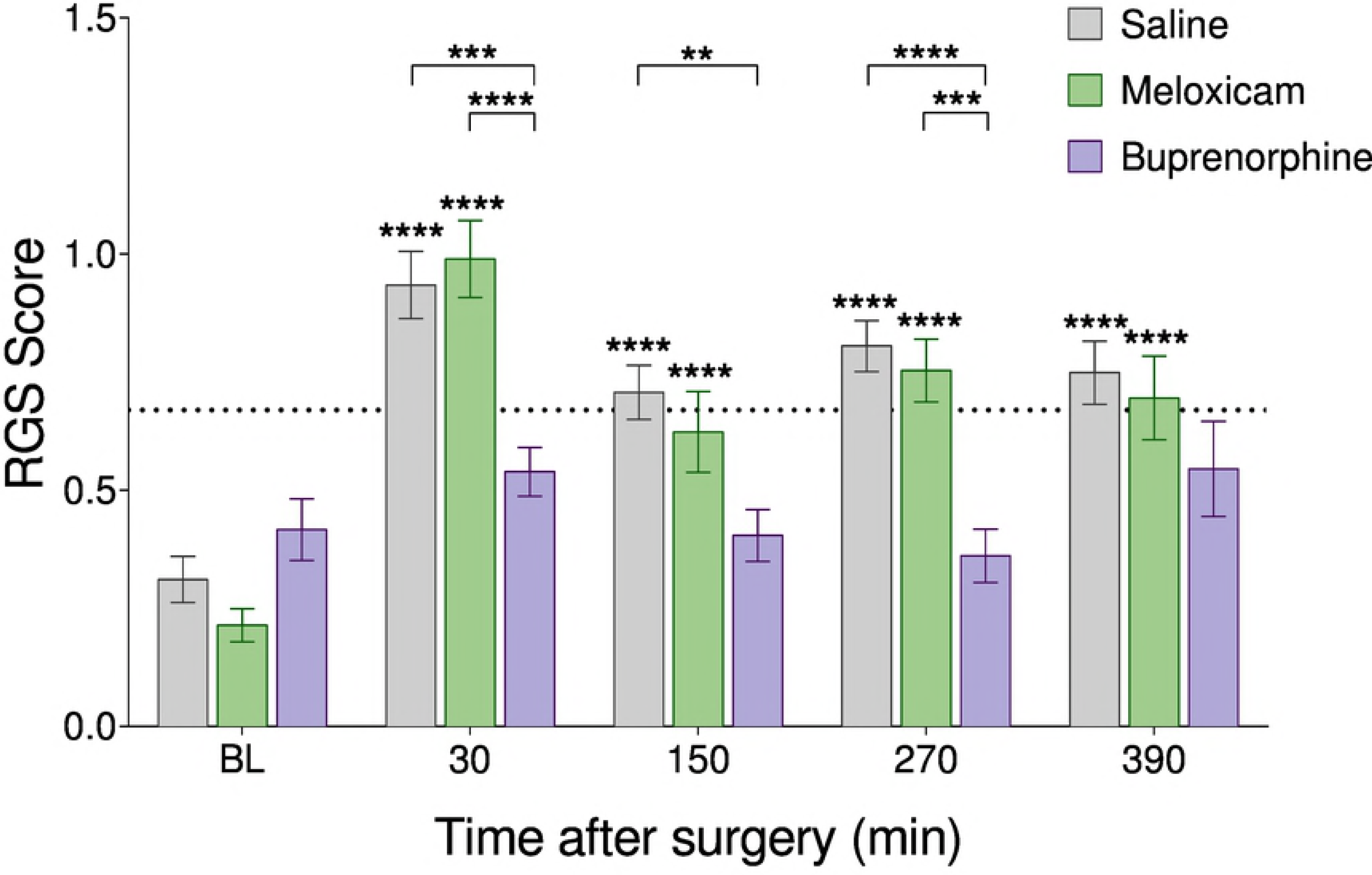
Rat Grimace Scale scores are reduced by the administration of buprenorphine after laparotomy. Saline (n = 21), meloxicam (n = 16) and buprenorphine (n = 15) groups at all time points. Data presented as mean ± SEM. BL = baseline. Asterisks above bars indicate within group differences from baseline. Asterisks with brackets indicate differences between groups.

**Fig 2.**
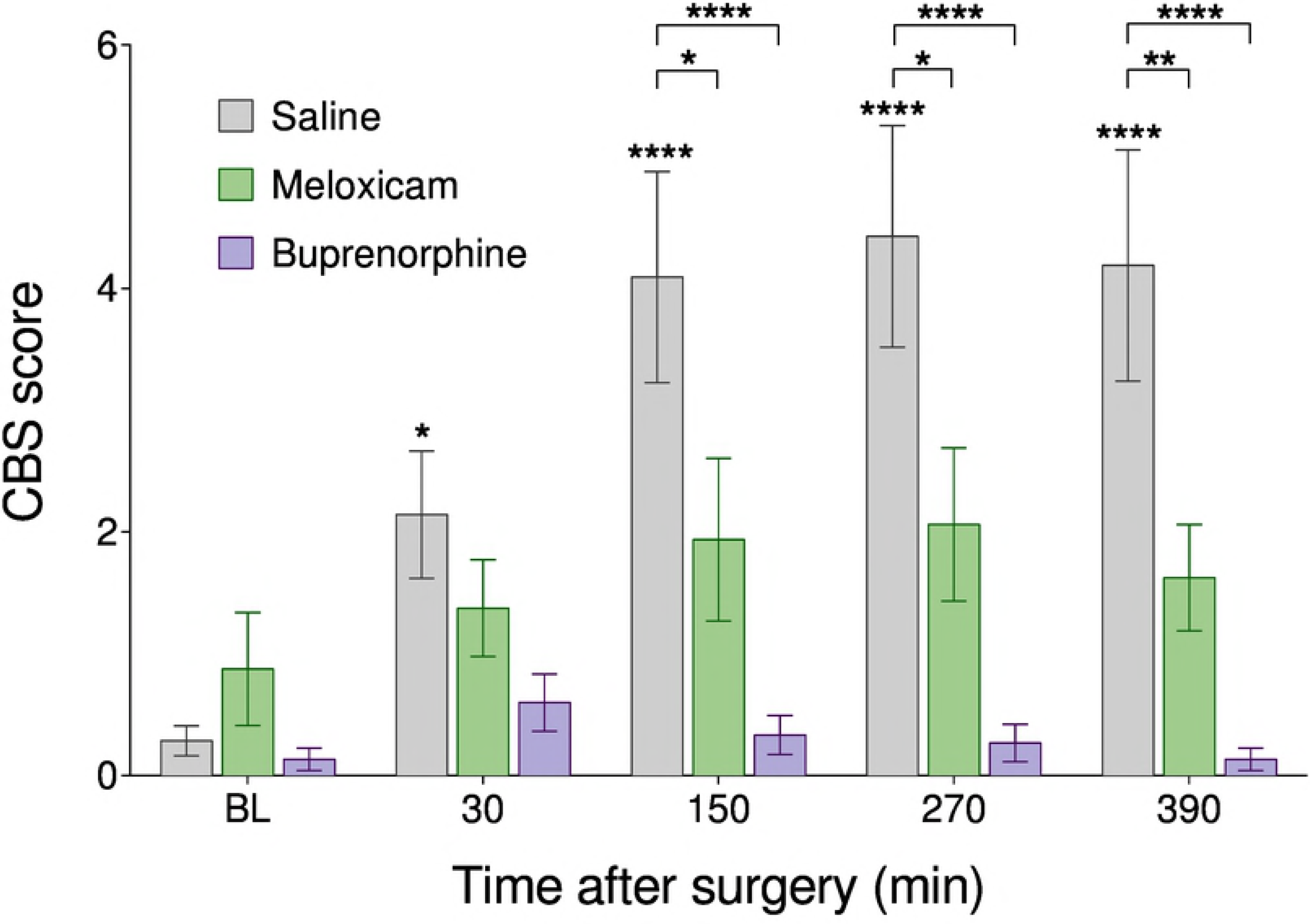
Composite behaviour scores (cumulative frequency of back-arching, writhing and staggering) is reduced by both meloxicam and buprenorphine following laparotomy. Saline (n = 21), meloxicam (n = 16) and buprenorphine (n = 15) groups at all time points. Data presented as mean ± SEM. BL = baseline. Asterisks stars above bars indicate within group differences from baseline. Asterisks with brackets indicate differences between groups.

### Between Group Comparisons

The RGS scores of saline- and meloxicam-treated animals were similar at all timepoints (p > 0.05; Table 2, Fig 1). Buprenorphine treated rats displayed lower RGS scores than saline treated rats at 30 minutes,150 minutes, and 270 minutes (p < 0.05) but not at baseline or 390 minutes (p > 0.05; Table 2, Figure 1). Buprenorphine treated animals also displayed lower RGS scores than meloxicam treated animals at 30 minutes and 270 minutes (p < 0.05) but not at baseline, 150 minutes, or 390 minutes (p > 0.05; Table 2, Fig 1).

**Table 2.** Between group comparisons of RGS and CBS scores at all timepoints.

| Comparison | RGS | CBS |
| --- | --- | --- |
|  | p-value [95% CI] | p-value [95% CI] |
| **Saline vs. Meloxicam** |  |  |
| BL | 0.55 [-0.12 to 0.31] | 0.76 [-2.5 to 1.4] |
| 30 min | 0.82 [-0.27 to 0.16] | 0.62 [-1.2 to 2.7] |
| 150 min | 0.64 [-0.13 to 0.30] | **0.026** [0.21 to 4.1] |
| 270 min | 0.84 [-0.17 to 0.27] | **0.013** [0.41 to 4.3] |
| 390 min | 0.83 [-0.16 to 0.27] | **0.0061** [0.61 to 4.5] |
| **Saline vs. Buprenorphine** |  |  |
| BL | 0.50 [-0.33 to 0.12] | 0.98 [-1.8 to 2.1] |
| 30 min | **0.0001** [0.17 to 0.62] | 0.16 [-0.45 to 3.5] |
| 150 min | **0.0042** [0.081 to 0.52] | **< 0.0001** [1.8 to 5.8] |
| 270 min | **< 0.0001** [0.22 to 0.67] | **< 0.0001** [2.2 to 6.2] |
| 390 min | 0.080 [-0.018 to 0.43] | **< 0.0001** [2.1 to 6.0] |
| **Meloxicam vs. Buprenorphine** |  |  |
| BL | 0.11 [-0.44 to 0.033] | 0.69 [-1.4 to 2.9] |
| 30 min | **< 0.0001** [0.21 to 0.69] | 0.66 [-1.3 to 2.9] |
| 150 min | 0.076 [-0.017 to 0.45] | 0.18 [-0.51 to 3.7] |
| 270 min | **0.0003** [0.16 to 0.63] | 0.11 [-0.32 to 3.9] |
| 390 min | 0.29 [-0.086 to 0.39] | 0.22 [-0.62 to 3.6] |

p-values and 95% confidence intervals of the differences are reported for each comparison.

Meloxicam treated animals displayed CBS scores significantly lower than saline treated animals at 150 minutes, 270 minutes, and 390 minutes (p < 0.05) but not baseline or 30 minutes (p > 0.05; Table 2, Fig 2). Similarly, buprenorphine treated animals also displayed CBS scores lower than saline treated animals at 150 minutes, 270 minutes, and 390 minutes (p < 0.05) but not at baseline or 30 minutes (p > 0.05; Table 2, Fig 2). Buprenorphine treated rats did not differ from meloxicam treated animals at any timepoint (p > 0.05; Table 2, Fig 2).

## Discussion

The need for better measures to assess ongoing pain in laboratory rodents has prompted the development of two behavioural pain scales: the RGS and CBS. Both pain scales have demonstrated construct validity as scores increase in response to common pain models and decrease with analgesic administration [7-10]. However, these two scales have never been directly compared and the assessment of their strengths and limitations, specifically their use to assess the efficacy of different analgesics, has been limited. The RGS and CBS have produced conflicting reports on the efficacy of NSAIDs to treat post-laparotomy pain in rats [8-10, 13]. Therefore, this study directly compared the two scales using a laparotomy pain model and two different classes of analgesics. We found that scores from both pain scales increased following laparotomy and were attenuated by buprenorphine. This is consistent with the large body of evidence that buprenorphine offers effective post-operative pain relief [13, 16, 17]. However, while CBS scores decreased with meloxicam administration, the RGS scores of the same rats did not decrease and were instead similar to saline treated rats. Therefore, this study provides the first direct evidence that these scales perform differently in their assessment of meloxicam, and perhaps other NSAIDs, as an effective analgesic following surgery. Together, these findings suggest that the CBS may be more sensitive at capturing analgesic responses than the RGS and that meloxicam may not provide suitable pain relief following laparotomy procedures.

This study echoes the discrepancy between previous studies using either the CBS or the RGS to assess the efficacy of NSAIDs [8-10, 13]. Specifically, low doses of NSAIDs reduced CBS but not RGS scores. Previous studies have shown that meloxicam at a dose of 1-2 mg/kg reduced CBS scores following laparotomy [9]. While the RGS has never previously been used to assess the efficacy of meloxicam following laparotomy, it has been used in conjunction with other pain models. It was reported that RGS scores decreased in response to meloxicam (2 mg/kg following a nerve root compression surgery) [18]. However, these scores were still elevated above the analgesic threshold, indicating that pain may still have been present, and were higher than the scores of control animals [18]. Interestingly, when rats were given meloxicam and buprenorphine together following an intra-plantar carrageenan model, these rats did not have lower RGS scores compared to rats that received buprenorphine alone [14].

Similar observations have also been reported with two other types of NSAIDs: ketoprofen and carprofen. Low doses of these NSAIDs have been repeatedly reported to reduce CBS behaviours, however, these same NSAIDs were reported as either ineffective at reducing RGS scores or a much higher dose was required [8-10, 13, 19, 20]. For example, ketoprofen at 5 mg/kg significantly reduced the frequency of CBS behaviours but a higher dose of 25-40 mg/kg was required to reduce RGS scores significantly [8, 13, 19, 20]. At high doses, it was reported that ketoprofen was as effective as 0.8 mg/kg of morphine at reducing RGS scores [20], however, it has also been reported that gastrointestinal side effects occur at doses as low as 5 mg/kg [13, 21, 22].

Interestingly, this requirement for high doses of NSAIDs to reduce grimace scale scores has also been observed in mice [23-25]. However, while 20 mg/kg of meloxicam was sufficient to reduce Mouse Grimace Scale (MGS) scores after a vasectomy in CD-1 mice, the same dose did not reduce MGS scores after a laparotomy in BALB/c mice even though meloxicam administration markedly reduced inflammation [23, 25]. The authors have suggested that this may have been related to differences in models or strains used [25]. Perhaps this could explain why a reduction in RGS score was observed after a nerve root compression surgery but not after a laparotomy [13, 18]. This highlights the need to study analgesic efficacy in different models and strains.

Overall, our study and previous studies seems to suggest that the CBS is more sensitive at discriminating analgesic efficacy of low doses of NSAIDs than the RGS. It is unknown if pain is completely abolished when the CBS scores are low as an analgesic intervention score for this scale has not been derived. The efficacy of meloxicam in rats could be further investigated with additional pain assessment tools (e.g. burrowing, to assess if motivation to perform highly enriching behaviours can be restored with meloxicam; or conditioned place preference testing, to assess if meloxicam administration can be positively associated). Additionally, while our study used only one dose of meloxicam, higher doses should be assessed to investigate whether this may provide more effective analgesia or if other unwanted effects occur. This would aid in concluding if the RGS is truly less sensitive than the CBS and if meloxicam should be used as an analgesic in rats.

## Meloxicam vs Buprenorphine

Buprenorphine treatment reduced both CBS and RGS scores to baseline levels in this study and in others [9, 13, 14, 16]. Therefore, it seems that buprenorphine is the more effective analgesic. However, buprenorphine use has been known to produce unwanted behavioural side effects including increased activity and self-injurious and pica behaviour [17, 26]. This has resulted in the use of NSAIDs as a popular alternative. However, the results of this study and other studies report that pain scores do not reduce to baseline levels when low doses of meloxicam and other NSAIDs are used alone [8-10, 13, 18, 23, 24]. This suggests that low doses of NSAIDs alone are insufficient to completely abolish the pain experience and a multimodal approach should be considered.

## CBS vs RGS methodology

While the CBS appears to provide a more sensitive measure of NSAID analgesia, it has some limitations. The CBS has received criticism as being too labour intensive [13]. In our video assessments, we obtained a pain score over 10 minutes; however, it has been reported that accurate pain scores can be obtained in as little as 5 minutes [9]. Nonetheless, compared to the RGS, this method has not yet been validated for real time observation as opposed to video analysis [14]. Additionally, the data variability of the CBS scores observed in the current study suggests that this method is more suitable for a research setting where data is averaged, as opposed to assessing pain in individual animals to identify if analgesic intervention is necessary. Thus, this limits its use in a clinical setting. In contrast, validation of RGS real-time scoring and derivation of an analgesic intervention threshold increase the value of the RGS as a tool for clinical assessment [11, 14].

## Limitations

A limitation of this study was that only female rats were used. Additionally, the administration of analgesics and anesthetics may have affected the scores through other mechanisms unrelated to pain. Particularly, the 30 minute timepoint may be particularly sensitive in this study as residual effects of the recent isoflurane anesthesia may affect behaviour and facial expression. Further research is needed to understand how analgesic and anesthetic administration affects the scores of these pain scales in the absence of pain.

## Conclusions

This study provides direct evidence that the RGS and CBS differ with regards to evaluating the efficacy of meloxicam in a laparotomy pain model. This suggests that the CBS may be more sensitive than the RGS in discriminating analgesic efficacy. However, further studies are required to assess if a low dose of meloxicam is truly efficacious for pain treatment and if both the CBS and RGS are assessing similar aspects of pain. The continued development of these methods of assessing ongoing pain is crucial to improving pain research and laboratory animal welfare.

## Supporting information

**S1 Table. Between strain comparisons of RGS and CBS scores of Sprague-Dawley and Wistar rats within treatment groups**. Sprague-Dawley: saline: n = 11, meloxicam: n = 8, buprenorphine: n = 7; Wistar: saline: n = 10, meloxicam: n = 8, buprenorphine: n = 8. p values and 95% confidence intervals of the differences are reported for each timepoint.

